# Prothrombin prevents fatal T cell-dependent anemia during chronic virus infection of mice

**DOI:** 10.1101/2024.11.11.623028

**Authors:** Rachel Cantrell, H. Alex Feldman, Leah Rosenfeldt, Ayad Ali, Benjamin Gourley, Cassandra Sprague, Daniel Leino, Jeff Crosby, Alexey Revenko, Brett Monia, Stephen N. Waggoner, Joseph S. Palumbo

## Abstract

Thrombin promotes the proliferation and function of CD8+ T cells. To test if thrombin prevents exhaustion and sustains antiviral T cell activity during chronic viral infection, we depleted the thrombin-precursor prothrombin to 10% of normal levels in mice prior to infection with the clone 13 strain of lymphocytic choriomeningitis virus. Unexpectedly, prothrombin insufficiency resulted in 100% mortality after infection that was prevented by depletion of CD8+ T cells, suggesting that reduced availability of prothrombin enhances virus-induced immunopathology. Yet, the number, function, and apparent exhaustion of virus-specific T cells were measurably unaffected by prothrombin depletion. Histological analysis of the lung, heart, liver, kidney, spleen, intestine, and brain did not reveal any evidence of hemorrhage or increased tissue damage in low prothrombin mice that could explain mortality. Viral loads were also similar in infected mice regardless of prothrombin levels. Instead, infection of prothrombin-depleted mice resulted in a severe, T cell-dependent anemia associated with increased hemolysis. Thus, thrombin plays an unexpected protective role in preventing hemolytic anemia during virus infection, with potential implications for patients who are using direct thrombin inhibitors as an anticoagulant therapy.

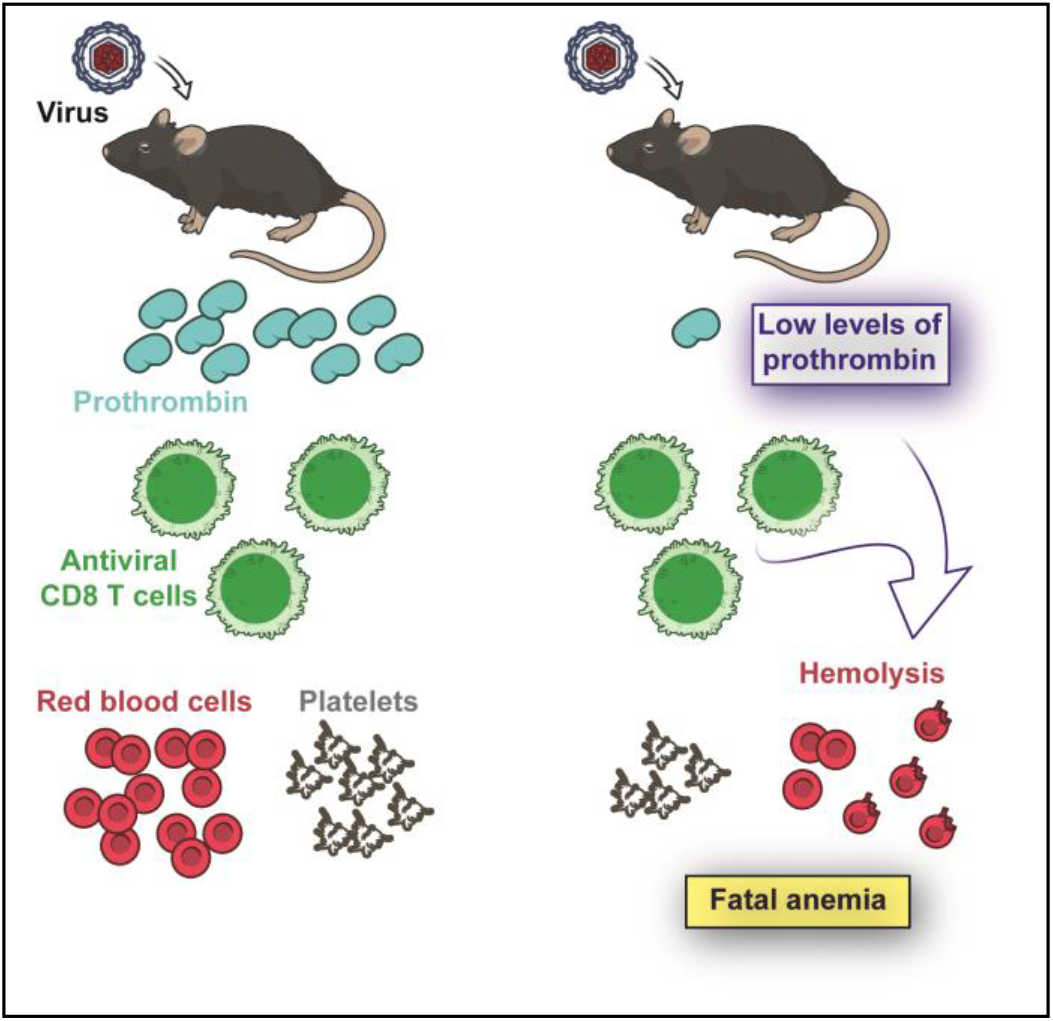

## Introduction

The generation of thrombin, the central hemostatic protease, is a ubiquitous feature of tissue injury (1). Prothrombin (FII) is converted to the active protease, thrombin (FIIa), through a series of proteolytic reactions termed the coagulation cascade. Coagulation can be initiated by the exposure of tissue factor (TF), which is abundantly expressed by cells in the subendothelial compartment (2). TF is the primary initiator of coagulation when there is disruption of vascular integrity. TF is also expressed by inflammatory activation of endothelial cells, as well as activated myeloid cells. Alternatively, coagulation can be initiated by the activation of factor XII, a protease that autoactivates when bound to negatively charged macromolecules such as DNA, which can be released at sites of tissue injury (3). Regardless of how the hemostatic cascade is initiated, thrombin activation is a key result of this process.

Thrombin is a promiscuous protease with at least 14 recognized substrates (1). Activated thrombin regulates hemostasis through proteolytic activation of both pro- and anti-coagulant proteases. Thrombin also regulates the activation and release of proteins involved in immune regulation, including IL-1α and TGFβ1 (1). In addition, thrombin directly regulates cellular processes by the activation of protease activated receptors (PARs), which are expressed by nearly all cell types, including platelets and multiple immune cells (4, 5). The potential for thrombin activation with traumatic tissue injury, inflammation, and infection make thrombin an important determinant of immune functions.

Recent evidence suggests that thrombin regulates T-cell responses (6-11). Human CD8+ T cells stimulated in the presence of thrombin exhibit elevated cytokine production and proliferation (6). Thrombin initiates Ca^2+^ flux and protein kinase C translocation to enhance early events in T cell activation (9-11), providing a possible mechanism by which thrombin can enhance T cell proliferation and functionality. Moreover, the thrombin-activated receptors, PAR-1 and PAR-4, are implicated in the regulation of CD4+ and CD8+ T cell function (11-15). Only PAR-1 has been studied in the context of T-cell responses to virus infection, with PAR-1-deficient mice exhibiting compromised antiviral CD8+ T cell function and viral control during acute infections with lymphocytic choriomeningitis virus (LCMV) (13). Thus, current evidence in mice and humans demonstrates the role of thrombin in supporting T cell function.

Based on these data, we assessed the role of thrombin in the maintenance of antiviral T cell function during chronic virus infection. High viral loads across multiple tissues after infection of mice with the clone 13 strain of LCMV promotes functional exhaustion and deletion of virus-specific T cells, resulting in a non-pathogenic chronic infection (16-18). Ablation of factors contributing to T cell exhaustion in this infectious context, including programmed cell-death 1 (PD-1) and natural killer (NK) cells, promotes a marked increase in the magnitude and function of antiviral T cells that contributes to elevated tissue damage and mortality (19, 20).

In contrast to the premise that thrombin supports T cell function, we find that reducing levels of prothrombin prior to LCMV infection results in uniformly fatal outcomes. This mortality was dependent on CD8^+^ T cells, but there was no evidence of elevated tissue pathology or measurable enhancement in the function of antiviral T cells. Instead, we identify a surprising role for prothrombin in preventing hemolysis and fatal anemia during chronic infection.

## Results

### Chronic virus infection becomes lethal in the setting of reduced prothrombin levels

To define the role of prothrombin (and thrombin) during chronic viral infection, we used an established prothrombin-specific antisense oligonucleotide (ASO) to reduce circulating prothrombin (FII) levels in C57BL/6 mice to ∼10% of normal levels (hereafter referred to as FII^Low^) (21-24). This achieved a concentration that roughly mimics the levels of prothrombin present in patients treated with high levels of warfarin (25). Control mice received the same dosing regimen of an oligonucleotide of similar chemistry with no homology in the mouse transcriptome. Intravenous inoculation of these mice with 2×10^6^ plaque-forming units of the clone 13 strain of lymphocytic choriomeningitis virus (LCMV clone 13) resulted in 100% mortality of prothrombin-depleted mice within two weeks of infection, while all control-treated mice survived (**Figure 1A**).

**Figure 1.**
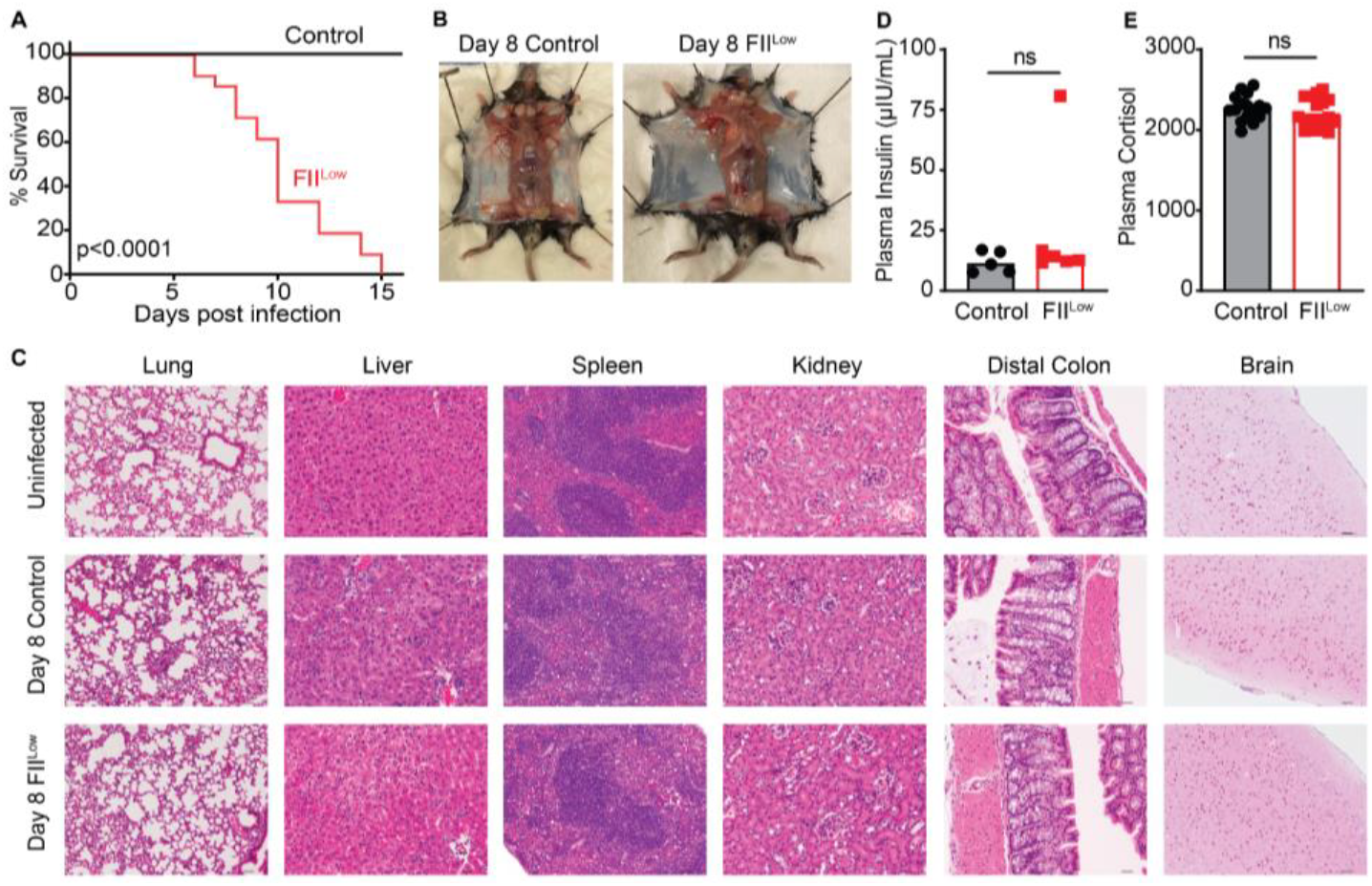
Prothrombin is required for survival of mice during chronic virus infection. **(A)** Survival of wild-type C57BL/6 mice that underwent pharmacologic depletion of prothrombin (FII^Low^, red squares) or control treatment (black circles) following intravenous infection with 2×10^6^ PFU of clone 13 LCMV (n=14-21/group). Absence of **(B)** subcutaneous hemorrhage and **(C)** representative H&E stained histological sections from lung (10x, 50μM scale, n=4-7), liver (20x, 25μM scale, n=12-17), spleen (10x, 50μM scale, n=8-13), kidney (20x, 25μM scale, n=4-7), distal colon (20x, 25μM scale, n=4-7), and the brain (10x, 50μM scale, n=8) of uninfected (n=4), control infected, and prothrombin-depleted infected mice on day 8 p.i. Levels of **(D)** insulin (n=5-6/group) and **(E)** cortisol (n=16/group) were measured in plasma on day 8 p.i. Mean ± SEM shown. Data are representative of results in at least two independent experimental replicates. Statistical significance was determined by Log-rank test for the survival curve, unpaired Student’s t-test or Mann-Whitney test, depending on normality of the data.

Given the role of thrombin in clotting (26) and the ability of arenavirus infections (such as LCMV) to cause lethal hemorrhage in susceptible hosts (27, 28), we assessed the possibility that significant hemorrhage is involved in the lethality of LCMV infection in prothrombin-depleted mice. However, extensive post-mortem analyses, including histological analyses of brain, lungs, heart, liver, spleen, kidney, and intestines, did not reveal any significant sites of hemorrhage in infected mice regardless of prothrombin levels (**Figure 1B**). Hemoccult tests for blood in the stool of infected mice were also uniformly negative (data not shown). Furthermore, intravenous administration of Evans Blue on day 6 of infection, 2 hours prior to euthanasia, revealed no significant vascular leak in control or low prothrombin mice that were infected with clone 13 LCMV (**Supplemental Figure 1**). Thus, there is no appreciable prothrombin-dependent increase in brain vasculature permeability.

In contrast to other models of lethal immunopathology during chronic infection with the clone 13 strain of LCMV, we did not observe enhancement of pathological damage to lungs (20, 29-31), liver (31-33), heart (19), kidney (34), brain (35), or intestines (36, 37) linked to fatal outcomes of infection in prothrombin-depleted animals (**Figure 1C, Supplemental Figure 2A-B**). Virus-infected, prothrombin-depleted mice also displayed similar plasma cortisol and insulin levels as control infected animals, decreasing the likelihood of adrenal insufficiency or acute metabolic abnormalities as the cause of death (**Figure 1D-E**). These results indicate that prothrombin protects mice from lethal outcomes of chronic virus infection in a manner distinct from known bleeding or organ damage pathologies previously linked to this virus.

**Figure 2.**
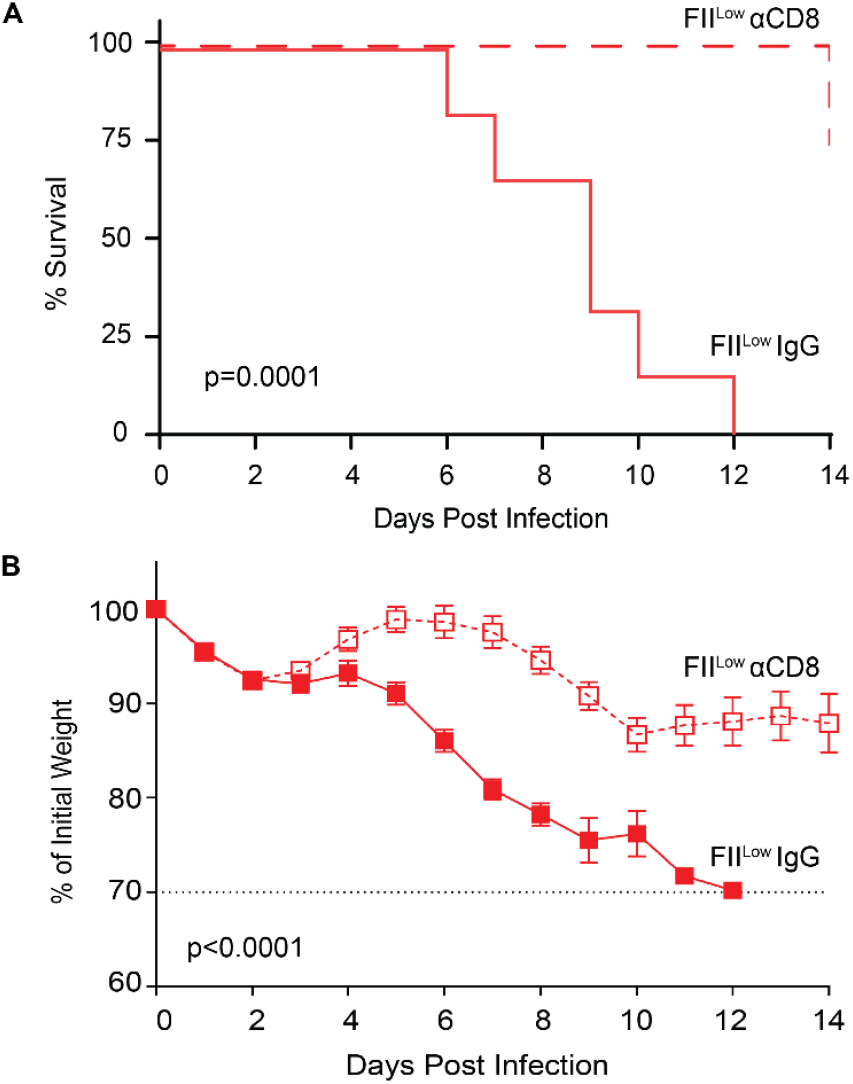
Prothrombin prevents fatal CD8+ T cell-mediated immunopathology. Prothrombin-depleted mice were injected with a CD8-depleting antibody (FII^Low^ αCD8, dashed red line, open red squares) or a control nonimmune IgG (FII^Low^ IgG, solid red line, solid red squares) prior to i.v. infection with 2×10^6^ PFU of the clone 13 strain of LCMV. Shown are **(A)** survival (n=6-8) and **(B)** weight loss (mean ± SEM, n=16-18) of these animals over time. Blue dashed line represents humane endpoint relative to weight loss, where all mice succumbed prior to reaching this limit. Statistical significance was determined by Log-rank test for survival curve. Stars on weight loss curve determined by unpaired t-test or Mann-Whitney test, depending on normality of data on each day post-infection, where *p=0.028, ***p=0.0002-0.0004, and ****p<0.0001.

### Prothrombin prevents CD8^+^ T cell-driven mortality during virus infection

As LCMV is a non-cytopathic virus, mortality is typically a result of T cell-driven immunopathology (38). Correspondingly, depletion of CD8^+^ T cells prior to infection of prothrombin-depleted mice prevented mortality (**Figure 2A**). CD8^+^ T cell depletion also abrogated weight loss following LCMV infection (**Figure 2B**). Thus, low prothrombin levels prompt fatal CD8^+^ T cell-driven immunopathology during chronic LCMV infection.

### Thrombin promotes in vitro survival, proliferation, and function of CD8+ T cells

Given that depletion of prothrombin unleashed fatal CD8^+^ T cell-dependent activity in infected mice, we hypothesized that the active form of thrombin directly regulates CD8^+^ T cell biology. Past reports suggest that thrombin can enhance T cell receptor signaling, cytoskeletal polarization, and cytokine production in human CD8^+^ T cells (6-11). We made similar observations in cultured mouse CD8^+^ T cells, with titrated depletion of thrombin from *in vitro* cultures of anti-CD3/anti-CD28 antibody-stimulated CD8^+^ T cells resulting in progressively worse T cell survival (**Figure 3A**), IFN-γ production (**Figure 3B**), and proliferation (**Figure 3C**). Thus, thrombin can bolster CD8^+^ T cell responses *in vitro*.

**Figure 3.**
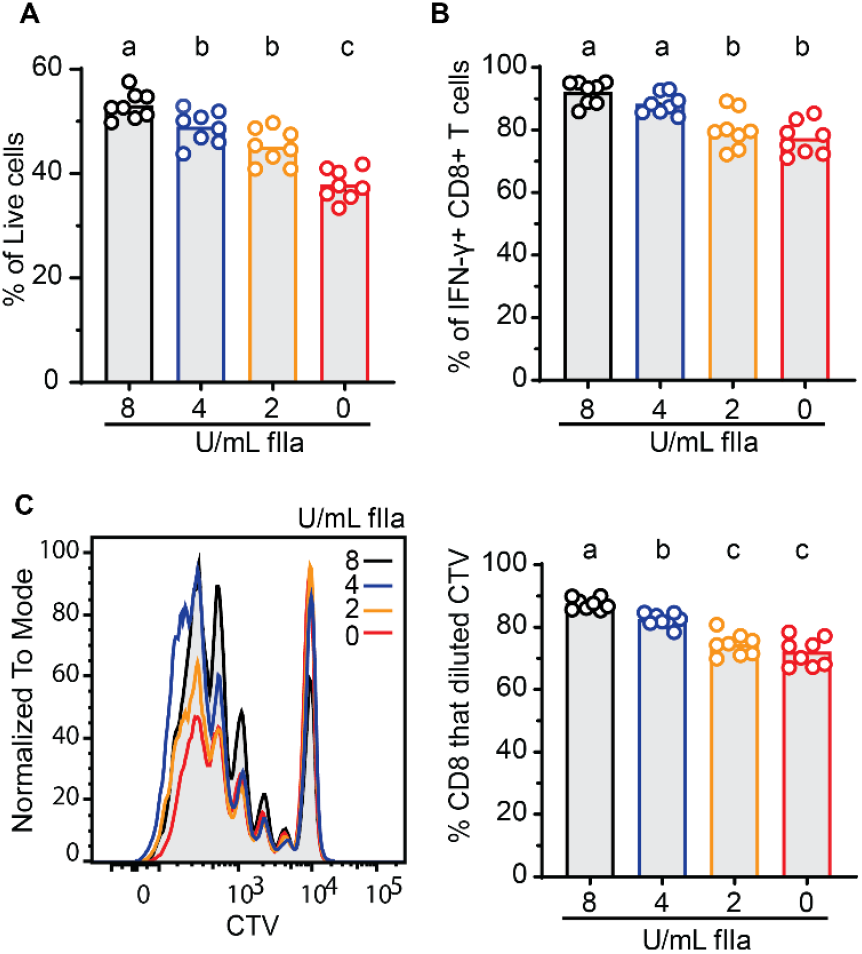
Thrombin promotes CD8+ T cell survival, proliferation, and function *in vitro*. Wild type C57BL/6 spleen CD8+ T cells were isolated, labeled with CellTraceViolet (CTV) and stimulated with αCD3+αCD28 antibodies for 72 hours in the presence (2, 4, 8 U/mL) or absence (0 U/mL) of thrombin. Flow cytometry was used to determine proportions (mean ± SEM) of **(A)** live, **(B)** IFN-γ+, and **(C)** proliferated CD8+ T cells in each condition. Data represents 8 technical replicates from pooled spleens of 4 mice. Data are representative of at least six independent experimental replicates. Means followed by a common letter are not significantly different by ordinary one-way ANOVA, followed by Tukey’s multiple comparisons test, with single pooled variance

### Prothrombin levels do not measurably affect antiviral CD8+ T cell responses in vivo

Given the supportive role of thrombin for CD8^+^ T cells *in vitro* and the fatal CD8^+^ T cell-dependent immunopathology in prothrombin-depleted mice (**Figures 1 and 3**), we assessed the consequence of prothrombin depletion on the magnitude and function of antiviral T cell responses *in vivo* (**Figure 4**). The proportions (representative gating shown in Figure 4A) and numbers of activated (CD44^hi^ CD43^+^) CD8^+^ (**Figure 4B**) and CD4+ (**Figure 4C**) T cell responses in the spleen at day 8 of infection were similar in prothrombin-depleted (FII^Low^) and control mice. There were no significant differences in the proportion or number of splenic CD8+ T cells that produced IFN-γ in response to *ex vivo* restimulation with the LCMV-derived GP_33-41_, GP_34-41_, GP_276-286_, NP_205-212_, or NP_396-404_ peptides between prothrombin-depleted and control mice, with the expected hierarchies among these responses during LCMV clone 13 infection (**Figure 4D**) (17). Likewise, the magnitudes of LCMV GP_64-81_-specific IFN-γ+ CD4+ T cell responses were similar in the spleens of prothrombin-depleted and control mice (**Figure 4E**). The cytolytic function of virus-specific CD8+ T cells against LCMV GP_33-41_ and NP_396-404_ peptide-coated target cells in vivo was similar in prothrombin-depleted and control mice at day 5 of infection (**Figure 4F**). Using immunohistochemistry, we observed no significant differences in the infiltration of CD8α+ T cells into the brains or livers of infected mice with normal or low levels of prothrombin (**Supplemental Figure 2C**). The relatively unchanged magnitude of antiviral T cell responses in prothrombin-depleted and control mice contrasts sharply with other contexts of fatal immunopathology during LCMV clone 13 infection, where fatal outcomes were associated with substantial (2.5-to 12.5-fold) enlarged pools of antiviral T cells (20, 31, 34, 39).

**Figure 4.**
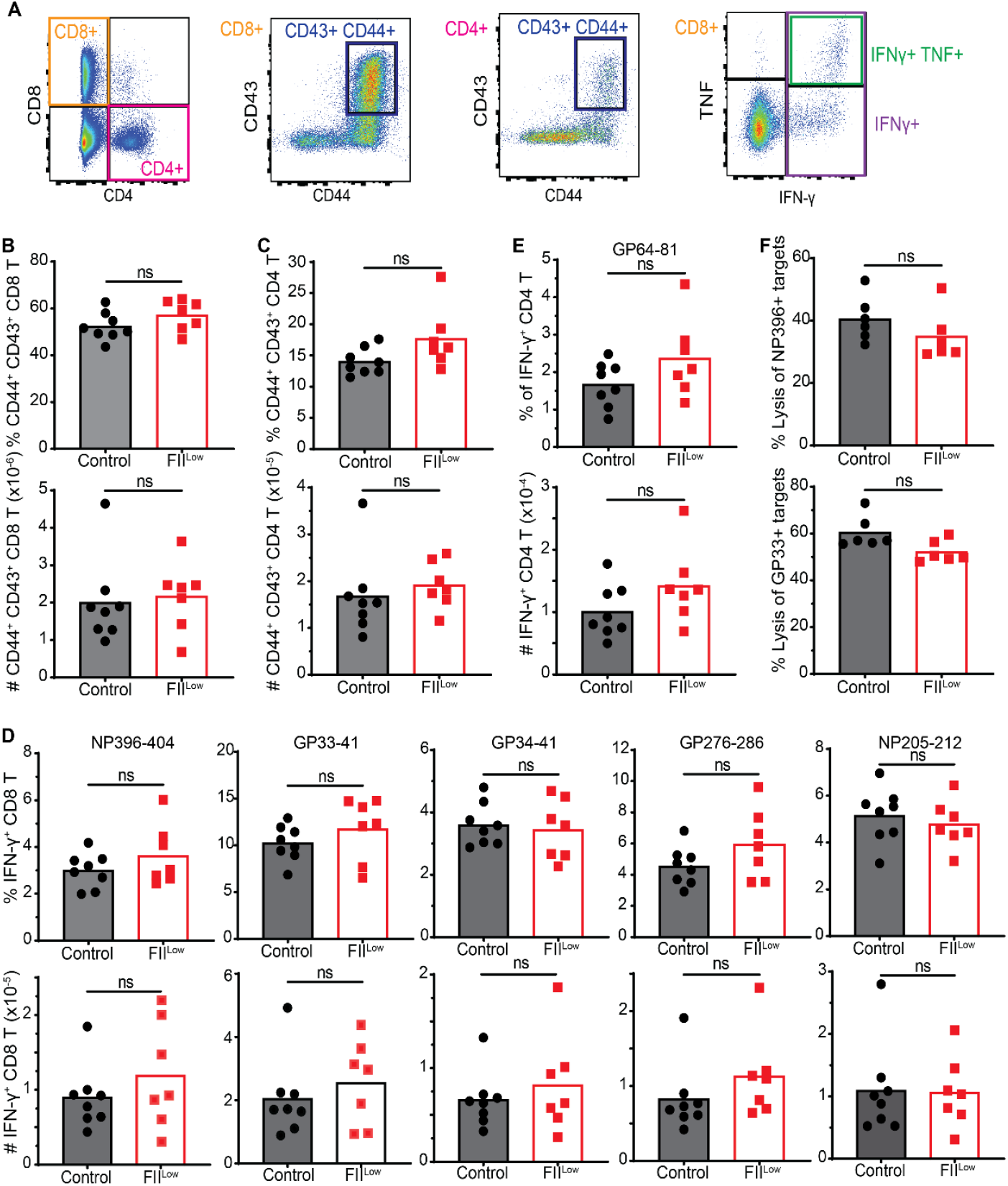
Magnitude of antiviral T cell responses is independent of prothrombin. Groups of C57BL/6 mice were treated with ASO to deplete prothrombin (FII^Low^) or control ASO (Control) prior to i.v. infection with 2×10^6^ PFU of the clone 13 strain of LCMV. **(A-E)** On day 8 of infection, antiviral T cell responses were analyzed in splenocytes (n=7-8 mice/group) following in vitro restimulation with LCMV-derived peptides. **(A)** Representative gating of CD4+ and CD8+ T cells among live, singlet lymphocytes as well as subsequent gating of CD44+CD43+, IFN-*γ*^+^, IFN-*γ*^+^TNF^+^ T cells is shown. The proportions and absolute numbers of activated CD44+CD43+ **(B)** CD8+ and **(C)** CD4+ T cells, as well as **(D)** IFN-*γ*+ CD8+ T cells and **(E)** IFN-*γ*+ CD4+ T cells stimulated with the noted viral peptides are presented. **(F)** For measurement of in vivo CTL function, splenocytes from uninfected mice were labeled with three different concentrations of CFSE, pulsed with viral-specific peptides (no peptide, GP_33-41_, or NP_396-404_), and intravenously transferred in a 1:1:1 ratio into prothrombin-depleted (FII^Low^) or control mice five days after infection (n=6 mice/group). Representative histogram of CFSE-labeled target cell recovery from a recipient mouse and mean (± SEM) calculated lysis of GP_33-41_^-^ or NP_396-404_-labeled target cells relative to unlabeled controls is plotted. Statistical significance was determined by unpaired Student’s t-test.

While the magnitudes of T-cell responses were similar in prothrombin-depleted and control mice after infection, we posited that functional exhaustion may be less efficient in the context of low prothrombin. However, expression levels of the key checkpoint receptor programmed cell-death 1 (PD1) were similar on CD8+ T cells in FII^Low^ and control mice (**Figure 5A**). Moreover, polyfunctionality as a measure of the capacity of virus-specific T cells to co-produce IFN-γ and TNF (**Figure 5B**) was similar in control and prothrombin low mice for all of the viral epitope-specific T cell responses tested (**Figure 5C**). Sera levels of IFN-γ and TNF were also similar in both groups of infected mice (Figure 5D) and substantially (20-to 100-fold) lower than those documented in the prior reports of lethal LCMV-induced immunopathology (19, 20). Finally, spleen and liver viral loads at day 8 of infection were similar in mice regardless of prothrombin levels, indicating similar capacity for T-cell constraint of virus replication (**Figure 5E**). Altogether, these results suggest that prothrombin paradoxically prevents T cell-driven pathology during chronic virus infection without altering the number or function of virus-specific T cells.

**Figure 5.**
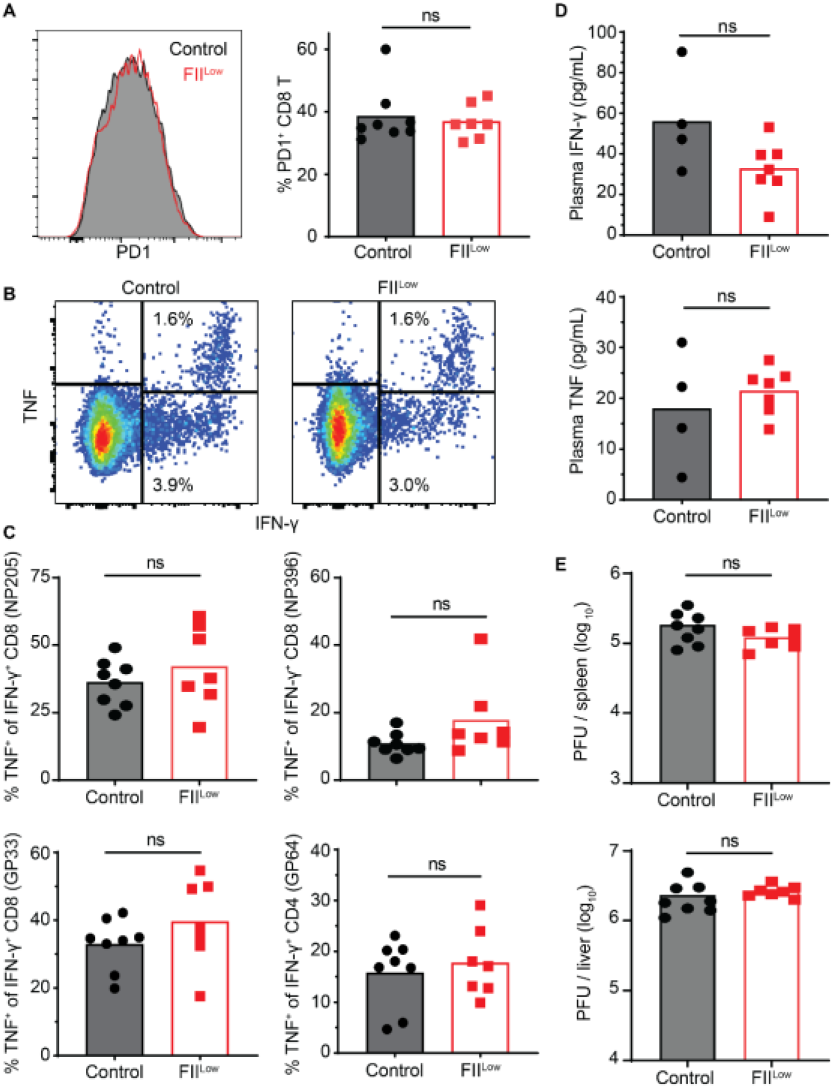
Prothrombin does not affect T cell exhaustion, function, or viral clearance. Further analysis of T cell function following i.v. infection of 2×10^6^ clone 13 LCMV into mice treated with FII ASO (FII^Low^) or Control ASO (Control) was performed and shown is **(A)** the proportion of CD8+ T cells expressing PD-1, with representative histogram of PD-1 expression levels in each group of mice (black line with grey fill=control, red line=FII^Low^). (**B**) Representative gating of IFN-*γ*^+^/TNF^+^ double-positive versus IFN-*γ*^+^ single-positive CD8 T cells stimulated with NP205 peptide. **(C)** Proportions of IFN-*γ*^+^ CD8+ (NP205, NP396, or GP33 stimulated) and CD4+ (GP64 stimulated) T cells that co-express TNF^+^ as a marker of functional exhaustion (n=7-8 mice/group). **(D)** Multiplex ELISA measurement of IFN-γ and TNF in plasma (n=4-7mice/group). **(E)** Plaque assay determination of viral titers in liver and spleen on day 8 of infection (n=7-8 mice/group). Statistical significance was determined by unpaired Student’s t-test.

### Prothrombin prevents severe anemia following chronic virus infection

Severe LCMV infection can result in anemia and thrombocytopenia (29, 30, 40-42). Therefore, we evaluated complete blood counts (CBC) on day 8 pos t infection. CBCs revealed that depletion of prothrombin was associated with a significant reduction in hemoglobin, hematocrit, and platelet counts as compared to control-infected mice (**Figure 6A**). Consistent with Figures 4 and 5, white blood cells counts were similar between the two groups of mice (**Figure 6A**). Although platelet count was reduced in the infected FII^Low^ mice compared to controls, sera PF4 levels as a marker of platelet activity were not significantly different between the two groups (**Figure 6B**). Thus, thrombocytopenia is not likely to be a significant contributor to mortality in this model, since the remaining platelets in mice with low prothrombin levels remain functional. In contrast, the degree of anemia observed in prothrombin-depleted mice was compatible with a potential cause of mortality (43). We also investigated the histology of the bone marrow (44, 45), but saw no prothrombin-dependent changes in hemophagocytosis, cellularity, or architecture (**Figure 6C**). Importantly, the lack of apparent hemorrhage in prothrombin-depleted mice suggests that anemia is likely independent of hemorrhage.

**Figure 6.**
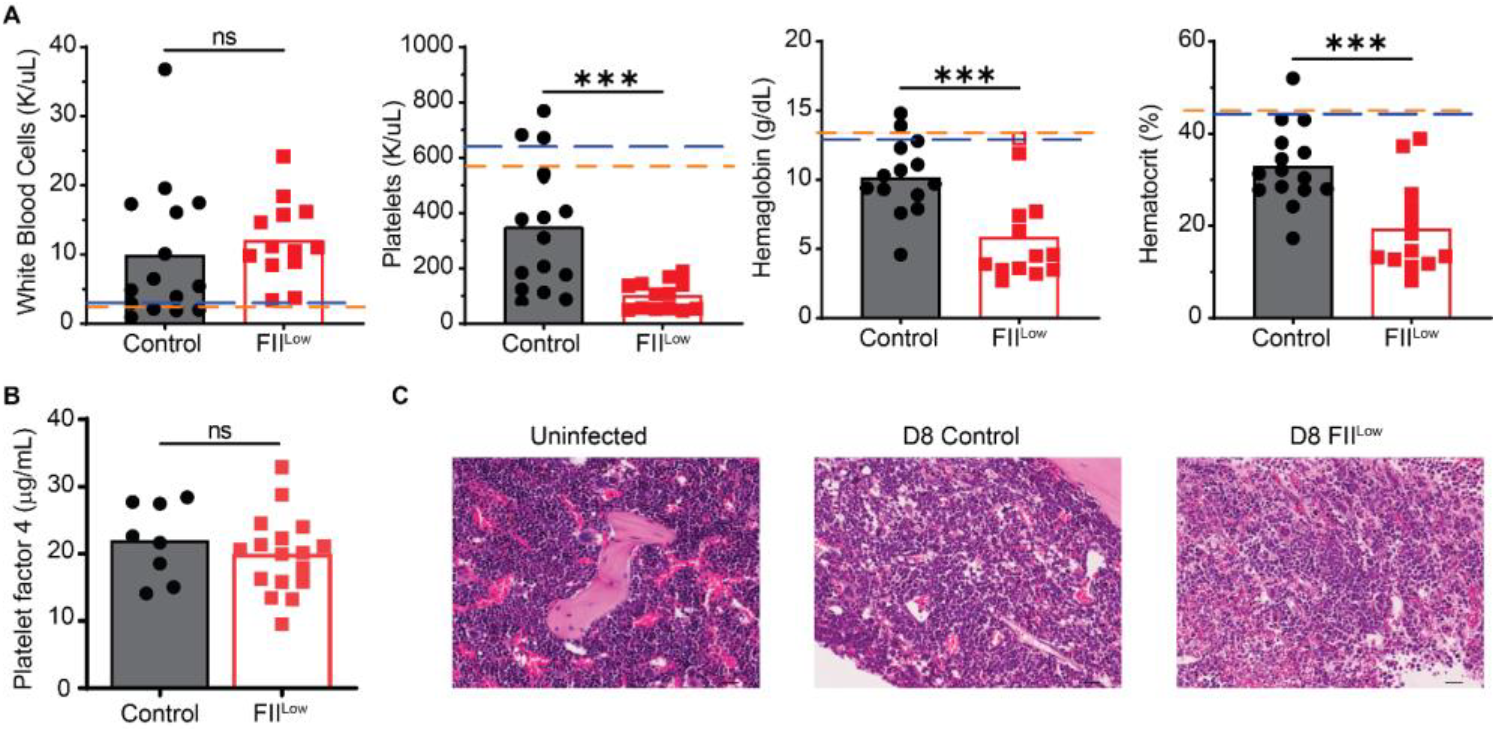
Prothrombin prevents severe anemia following chronic virus infection. Groups of C57BL/6 mice (n=13-15) were treated with ASO to deplete prothrombin (FII^Low^) or control ASO (Control) prior to i.v. infection with 2×10^6^ PFU of the clone 13 strain of LCMV. **(A)** On day 8 of infection, complete blood count analysis was performed to determine white blood cell and platelet counts as well as hemoglobin and hematocrit levels (n=13-15). Dashed lines represent mean values measured in uninfected control (blue line) and FII^Low^ (Orange line) mice (n=4). **(B)** PF4 ELISA was performed using plasma from these mice. **(C)** Femurs were harvested on 8 days p.i. and H&E staining was performed, with representative results from 4-6 mice/group shown. CBC analysis was repeated in two independent experiments. Statistical significance was determined by unpaired t-test or Mann-Whitney test, based on normality of data.

### Prothrombin ameliorates hemolysis and CD8+ T cell-dependent anemia

Analysis of plasma taken from control and prothrombin-depleted mice on day 8 of infection revealed that prothrombin depletion results in increased LDH and reduced haptoglobin (**Figure 7A**). These measures suggest that intravascular hemolysis is elevated in the setting of reduced prothrombin availability during chronic virus infection. Critically, depletion of CD8^+^ T cells in prothrombin-depleted mice restored the hemoglobin, platelets, and hematocrit measures to those observed in control animals (**Figure 7B**). T-cell derived IFN-*γ* can drive anemia during LCMV infection (40, 46, 47), yet neutralization of IFN-*γ* in infected low prothrombin did not prevent low hemoglobin, platelets, and hematocrit measures (**Supplemental Figure 3**). These data implicate prothrombin in prevention of an IFN-*γ*-independent, CD8+ T cell-driven hemolytic anemia during chronic LCMV infection.

**Figure 7.**
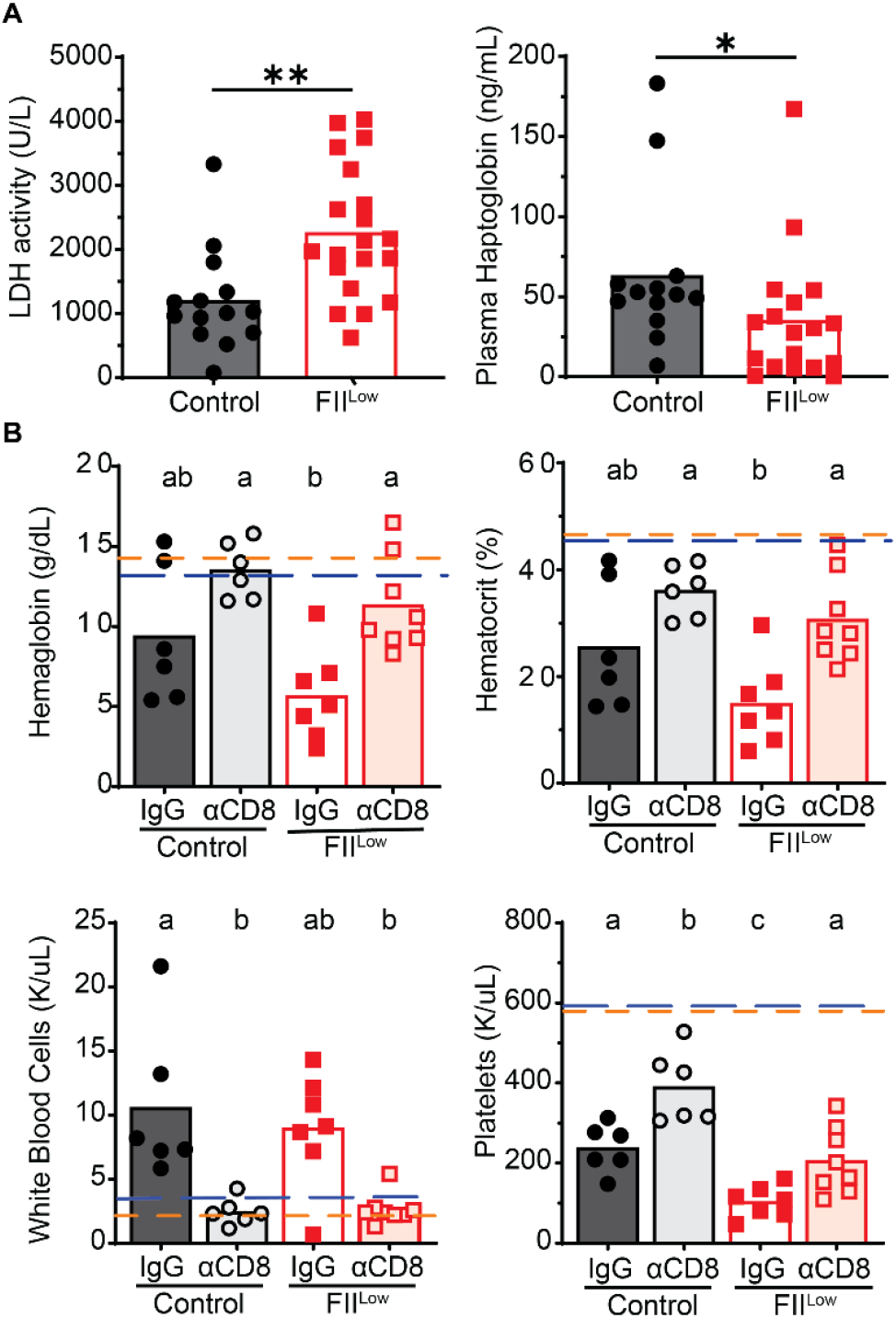
Prothrombin prevents CD8+ T cell-dependent anemia. Plasma was obtained on day 8 of infection from C57BL/6 mice treated with control ASO (Control) or ASO to deplete prothrombin (FII^Low^) prior to i.v. infection with 2×10^6^ PFU of the clone 13 strain of LCMV. **(A)** Plasma was used to assess LDH activity and haptoglobin levels (n=13-20). **(B)** Some groups of mice (n=6-8) were treated with anti-CD8 antibody (α) or IgG control prior to infection. Hemoglobin, hematocrit, white blood cell and platelet counts were assessed. Dashed lines represent mean values measured in uninfected control (blue line) and FII^Low^ (Orange line) mice (n=4). CBC data represents data from two independent experiments. Statistical significance was determined by Mann-Whitney test. Means followed by a common letter are not significantly different as determined by Kruskal-Wallis followed by Dunn’s multiple comparison test.

Although CD8 T cells were necessary for anemia, attempts to show that low prothrombin availability alters CD8 T cells in a manner sufficient to induce anemia were less conclusive. Adoptive transfer of 100,000 splenic CD8+ T cells from infected donor mice with low levels of prothrombin (FII^Low^) was insufficient to restore anemia in infected, CD8-depleted recipient mice to the levels observed in CD8-sufficient FII^Low^ mice (**Supplemental Figure 4**). Technical difficulties with this schema, including timing, number, and trafficking of donor T cells, preclude determination of whether CD8 T cell could be sufficient to drive anemia. However, an intermediate anemia phenotype was observed in T cell reconstituted FII^Low^ mice relative to T cell reconstituted control mice, regardless of whether donor T cells came from low or normal prothrombin settings (**Supplemental Figure 4**). This observation, coupled with the data in Figure 3, suggests that CD8 T cells may rapidly adapt to environmental levels of prothrombin and provoke anemia in synergy with complex infection- and prothrombin-driven factors.

## DISCUSSION

We demonstrate an unexpected requirement for prothrombin in survival of mice during chronic arenavirus infection. Decreasing prothrombin levels prior to infection resulted in the induction of severe anemia, thrombocytopenia, hemolysis, and death that were dependent upon CD8+ T cells. However, this enhanced T cell-driven immunopathology was uncoupled from any measurable increase in the number, function, or exhaustion of antiviral T cells. Moreover, anemia in the context of low prothrombin levels was not impacted by neutralization of IFN-*γ*. Thus, thrombin restrains the pathogenicity of antiviral CD8+ T cell responses in an unconventional manner to limit hemolytic anemia during chronic virus infection.

Since LCMV is a non-cytopathic virus, morbidity and mortality during infection are predominately caused by immunopathology. Dysregulated antiviral T cell responses during LCMV infection can provoke cytokine storm, organ damage, hemorrhage, and autoimmunity (19, 20, 30-35, 47, 48). This infection model has been used to identify multiple translationally relevant mechanisms that prevent these immunopathologic outcomes (38), including checkpoint receptors and T cell exhaustion (17-19, 33, 49). Interfering with these immunoregulatory mechanisms results in fatal pulmonary (20, 29-31), hepatic (31-33), cardiac (19), renal (34), intestinal (36, 37), vascular (28, 29, 50), splenic (42), pancreati c (51), or neurological (35) sequelae in an otherwise non-lethal infectious context. Here we identify prothrombin as a previously undescribed checkpoint preventing fatal immunopathology. Intriguingly, none of the established organ pathologies of overexuberant T cell responses during LCMV infection we analyzed were measurably exacerbated by the depletion of prothrombin. Moreover, there was no evidence of cytokine storm or failure of major organ systems in prothrombin-depleted mice. These results suggest that the protective effects of thrombin during LCMV do not involve prevention of common virus-induced immunopathological sequelae.

Instead, depletion of prothrombin prior to LCMV infection of mice resulted in severe anemia and thrombocytopenia that was dependent on the presence of CD8+ T cells. The observed thrombocytopenia is an unlikely cause of the observed mortality. First, the low platelet counts associated with reduced prothrombin availability (mean 100,000 platelets/µL) markedly exceed the levels (<20,000 platelets/µL) associated with hemorrhagic disease in LCMV-infected, platelet-depleted mice (28). Second, we find that the platelets in mice with low prothrombin levels remain functional. Given the role of thromb in in hemostasis and the hemorrhagic phenotype observed in some mice with severe thrombocytopenia during LCMV infection (28, 29, 32), we first suspected that increased bleeding in low prothrombin mice would be associated with this anemia. However, there was no evidence of hemorrhage in low prothrombin mice at any time point after infection when compared to published studies of LCMV-induced hemorrhage (28) or our own experience in mice with hemostasis deficiencies (52, 53). There was also no evidence of occult blood loss in the gastrointestinal tract or increased vascular leakage in the brain.

Other reported causes of anemia in LCMV infection include bone marrow failure and exaggerated hemophagocytosis (41, 47). Our pathology and flow cytometry-based characterizations of spleen and bone marrow in infected mice did not reveal evidence that hemophagocytosis is exacerbated or that erythropoiesis is impaired when there is limited availability of prothrombin. In the absence of overt mechanisms causing increased bleeding or failed generation of red blood cells during infection of low-prothrombin mice, we hypothesized that thrombin may subvert red blood cell destruction (hemolysis) to prevent fatal anemia during infection. Of note, CD4 T cells sustain CD8 T cell function during chronic LCMV infection and could be involved in this anemia, yet determination of an independent role for CD4 T cells would be difficult since CD4 depletion profoundly affects the responses of all T cell subsets in this model system (18, 54).

Indeed, elevated plasma LDH activity and low plasma haptoglobin levels in LCMV-infected, prothrombin-depleted mice indicate an increased severity of hemolysis in these animals. Anemia in these mice was dependent on CD8^+^ T cells, suggesting that altered T cell function in the setting of low prothrombin levels results in increased hemolysis. However, expression of anemia-promoting cytokines such as IFN-*γ* was relatively normal in mice with low prothrombin levels, and neutralization of IFN-*γ* did not prevent induction of anemia in FII^Low^ mice. Infections with LCMV are reported to induce autoimmune hemolysis resulting in mild anemia (55-58). Virus-induced autoimmune hemolytic anemia can be caused by autoantibodies or by direct activities of T cell, depending on the strain of mouse that is infected (28, 41, 55-60).

Therefore, the CD8^+^ T cell-dependent anemia in low prothrombin animals after infection could be a direct result of aggravated T cell functionality or an indirect regulation of a distinct red blood cell lysis mechanism (i.e. autoantibody driven) by CD8^+^ T cells. Physiological levels of prothrombin may prevent this hemolytic anemia in several ways, including direct regulation of T cell function (6, 7, 11). Of note, the number, function, and exhaustion of antiviral T cells all appeared similar between prothrombin-depleted and control mice. This finding uncouples prothrombin from conventional mechanisms of enhanced immunopathology during LCMV due to exaggerated T cell responses (19, 20, 30, 31, 33-35).

CD8 T cells exposed to low levels of prothrombin during activation did not appear to be sufficient to induce anemia following adoptive transfer into CD8-deficient hosts, although this experiment is technically challenging. Thus, an alteration of T cell function and/or an indirect regulation of downstream mechanisms that limit CD8^+^ T cell promotion of hemolytic anemia remain possible. Such indirect mechanisms could include thrombin regulation of complement activation (61) or reticuloendothelial clearance of antibody-bound red blood cells (62). Thrombin is also implicated in immune regulation via activation of latent TGF-β (63). Neutralization of TGF-β or ablation of TGF-β-receptor signaling in T cells modestly increases T cell responses against LCMV (64-67), in some cases resulting in immunopathology. As such, low levels of prothrombin in our system could promote increased pathology by reducing the availability of active TGF-β. Intriguingly, thrombin-mediated platelet activation via protease activated receptors results in a conformational change in αIIbβ3 integrin that increases affinity of the integrin for fibrin (68, 69). Loss of platelet β3 integrins results in severe anemia during LCMV infection (28). Thus, impaired thrombin-mediated platelet activation in the context of low prothrombin levels could promote anemia by impairing platelet-driven immunoregulatory mechanisms. Any of these CD8 T cell extraneous mechanisms could explain the low prothrombin environmental effects observed in the CD8 T cell transfer experiments. Though the precise mechanism remains to be explored, our data clearly point to a role for thrombin in preventing a CD8+ T cell dependent hemolytic anemia during chronic virus infection.

**T**ogether with previously published work, our findings add depth to the scientific understanding of immune- and hemostatic-mediated responses that are necessary to survive chronic virus infections. In the non-hemorrhagic virus infection model (70) used in this manuscript, we highlight a critical requirement of prothrombin in preventing anemia caused by CD8^+^ T cells that is independent of the magnitude and exhaustion of the antiviral T cell responses and associated production of IFN-*γ*. In hemorrhagic virus infections, severe disruption of platelet function (28) could potentially synergize with low levels of prothrombin to exacerbate bleeding. As many patients require anticoagulants for prevention of thrombosis, our findings raise the specter of potential adverse effects of using anti-thrombin therapies that may leave patients susceptible to infection-induced immunopathological anemia. Future studies will be needed to address the contribution of thrombin and anticoagulant therapies, which target thrombin or thrombin generation, in the survival and recovery of other viral infections in mice and humans.

## Materials and Methods

### Sex as a biological variable

One experiment was conducted using both male and female mice, with no differences observed between sexes in the induction of fatal immunopathology during LCMV infection in the context of low prothrombin levels. Thereafter, our experimental design focused on female mice with resulting findings expected to be relevant for all sexes.

### Mice

8-week-old C57BL/6J mice were purchased from The Jackson Laboratories. Mice between 8 to 20 weeks of age were routinely utilized in experiments. Mice were housed under barrier conditions and experiments were performed under ethical guidelines approved by the Institutional Animal Care and Use Committees of Cincinnati Children’s Hospital Medical Center. In most experiments, cage mates were randomly assigned to different experimental groups.

### Manipulation of prothrombin levels

Prothrombin (Factor II, FII) was lowered to about 10% of normal levels as previously described by weekly subcutaneous injections of 50 mg/kg prothrombin-specific antisense oligonucleotide (ASO) provided by Ionis Pharmaceuticals (21, 22). Control mice were treated in parallel with an oligonucleotide of similar chemistry with no homology in the murine transcriptome.

### In vivo CD8+ T cell depletion and IFN-γ neutralization

One day prior to infection, mice were injected retro-orbitally with 200 μg of *InVivo*MAb anti-mouse CD8α antibody (Clone: YTS 169.4; BioXCell) or control IgG2b isotype antibody (Clone: LTF-Z; BioXCell) (71). CD8+ depletion was confirmed by flow cytometry at the conclusion of each experiment. For experiments depleting IFN-*γ*, mice were infected with LCMV Clone 13 as described below. At Days 2 and 5 post-infection, mice were injected retro-orbitally with 200 μg of anti-mouse IFN-*γ* (Clone: XMG1.2; BioXCell) or control IgG1 isotype antibody (Clone: TNP6A7; BioXCell).

### Virus and infections

Stocks of the clone 13 strain of LCMV were generated using BHK21 cells, while viral titers in stocks, tissue samples, or blood were determined via plaque assay using Vero cells (72). One day after the third ASO injection, mice were infected retro-orbitally with 2×10^6^ plaque-forming units (PFU) of LCMV Clone 13. Daily body weight was recorded for the duration of the experiment. Mice experiencing greater than or equal to 30% weight loss were euthanized in accordance with Institutional Animal Care and Use Committee approved guidelines.

### In vitro T cell activation and exposure to thrombin

Spleens were harvested from C57BL/6J mice and placed in processing medium (Spinner’s Modification of Minimal Essential Media supplemented with 2% FBS, 1% L-glutamine and 2% penicillin/streptomycin, and 50 µM 2-Mercaptoethanol). Organs were dissociated and passed through a 70 µm filter. Red blood cells were lysed via exposure to ACK lysis buffer for one minute at room temperature before washing with processing media. CD8+ T cells were isolated with CD8+ T cell negative selection kit (Biolegend) following the manufacturer’s instructions with the following exception: MojoSort buffer was replaced with SMEM media supplemented with 10% FBS, 2mM EDTA, 1% L-glutamine and 2% penicillin/streptomycin and 50 µM 2-Mercaptoethanol so that bovine serum albumin was not introduced to the system. Isolated CD8+ T cells were stained with 3 µm/mL Cell Trace Violet diluted in 0.2% FBS in PBS for 20 minutes at 37°C. Free dye was removed by adding FBS and placing cells in 4°C for five minutes. Cells were plated on 1 µg/mL anti-CD3 antibody (Biolegend) pre-coated 96-well U-bottom plate at 2×10^5^ cells/well. Cells were further activated with 1 µg/mL of anti-CD28 antibody in the presence or absence of 2-8 U/mL of murine thrombin (Innovative Research, Novi, MI) at 37°C for 72 hours. Five hours before staining for flow cytometry cells were exposed to 1X protein transport inhibitors (BD) to block cytokine release.

### CD8+ Adoptive Transfer

Following prothrombin depletion, donor mice were infected with LCMV Clone 13 as outlined above. Meanwhile, we depleted CD8+ T cells in recipient mice one day before infection as described above. On day 6 post-infection, donor mice were euthanized and spleens processed as described previously. Next, we enriched CD8+ T cells using a mouse CD8α T Cell Isolation kit (Miltenyi Biotec) to >95% purity as verified by flow cytometry (data not shown). Afterwards, isolated CD8+ T cells were resuspended in HBSS at a concentration of 1×10^6^ cells/mL and 100 µL of cells was injected retro-orbitally into normal or FII^Low^ recipients on day 4 of infection. Mice were then monitored until euthanasia at day 8 post-infection.

### Histological analysis

Organs of interest were harvested from euthanized experimental mice and immediately placed in 10% formalin. After 24-hour fixation, organs were transferred into 70% ethanol until processing. Samples were sent through a long cycle in a Microm STP 120 Spin Tissue Processor (Thermo Fisher Scientific) prior to embedding in paraffin using a Tissue TEC (Sakura Finetek USA). Slides were cut to 5 µM thickness using a Leica RM2135 microtome (Leica Biosystems). Then, samples were deparaffinized in a vacuum oven and stained with hematoxylin and eosin following standard protocols. Images were taken on a brightfield microscope at 10X objective. When appropriate, slides were given to our pathologist for Batts-Ludwig scoring and evaluation of phenotypes while blinded to groupings. For immunohistochemistry staining, paraffin-embedded livers were sectioned and mounted on slides as described above. Slides were deparaffinized using xylene and rehydrated by incubating for 5 minutes each in different percentages of ethanol (100%, 95%, 70%, 30%) followed by 5 minutes wash with distilled water. Antigen retrieval was performed by heating the slides in pH 6.0 sodium citrate buffer in the microwave for 10 minutes. Slides were cooled down for 30 minutes at room temperature (RT) and washed with PBS. Endogenous peroxidase activity is quenched by incubating the sections in BLOXALL® Blocking Solution (Vector Laboratories) for 10 minutes and then washed with PBS for 10 minutes. Slides were incubated with PBS containing 10% normal goat serum for 60 minutes at RT for reducing the unwanted staining. For detection of CD8α, slides were incubated overnight at 4°C with CD8 alpha rabbit antibody (Clone: D4W2Z; Cell Signaling Technology) at a dilution of 1:200. Slides were washed 3 times with 0.1% PBST and then incubated with ImmPRESS Polymer Reagent (Vector Laboratories) for 30 minutes. Slides were washed with 0.1% PBST 3 times and incubated in peroxidase substrate solution (Vector Laboratories) for 5 minutes. Slides were counter stained with hematoxylin, dehydrated with ethanol and xylene and mounted with permount.

### Ex vivo organ preparation of spleen and bone marrow leukocytes

Mice were anesthetized by continuous isoflurane and euthanized via fatal blood draw of at least 300 μL into 10% citrate from the inferior vena cava. Afterwards, anesthetized mice were euthanized by cardiac dissection. Spleens were immediately removed from mice and placed on ice in 600 μL of RPMI-1640 (Cytiva) supplemented with 10% FBS, 1% Penicillin/Streptomycin, and 1% L-Glutamine before mechanical dissociation using glass microscope slides and filtering through a 70-micron filter to generate a single cell suspension. Splenocytes were then centrifuged for 5 minutes at 325x*g* before discarding the supernatant. Next, we added ACK lysis buffer (made in-house) and incubated splenocytes at 37°C for 5 minutes before repeating the earlier centrifugation step. Again, the supernatant was discarded before a final resuspension in 2 mL of complete RPMI-1640.

### Flow cytometry and intracellular cytokine staining

Following processing of organs, 100 μL of splenocytes were added to a U-bottom 96 well plate and pelleted at 325xg for 3 minutes before washing with FACS buffer (Hank’s Buffered Salt Solution without calcium or magnesium, 1% FBS, 2mM EDTA). Cells were then stained for 5 minutes with Zombie NIR Live/Dead (Biolegend) and washed twice with FACS buffer. Next, we incubated cells with Fc Shield (aCD16/CD32, Tonbo) for 5 minutes at 4°C before washing with FACS buffer. Cells were then stained with a combination of the following antibodies: CD4, CD8a, CD8b, CD41, CD43, CD44, CD49f, CD55, CD71, CD105, CD150, cKit. Afterwards, cells were centrifuged and washed once with FACS buffer before adding 100 μL of Cytofix (BD) and incubating for 4 minutes at 4°C. Finally, cells were centrifuged and washed twice with FACS buffer before resuspending in 200 μL of FACS buffer. All cells were counted using a hemocytometer and the trypan blue exclusion method.

To assay the activity of virus-specific T cells, 500 μg/mL of anti-CD3e antibody (Clone 145-2C11; Cytek Biosciences) was diluted in PBS to 10 μg/mL and added to a 96 well plate for positive control wells the night prior to each experiment. The day of the experiment, organs were processed as outlined above. 2 μg/mL of LCMV peptides (GP_33-41_, KAVYNFATC; GP_34-41_, AVYNFATC; GP_64-80_, GPDIYKGVYQFKSVEFD; GP_276-286_, SGVENPGGYCL; NP_205-212_, YTVKYPNL; or NP_396-404_, FQPQNGQFI) were mixed with 2μL of GolgiPlug (BD) per well and plated with 100μL of splenocytes for 4 hours at 37°C. Afterwards, cells underwent surface staining as described above. Next, cells were permeabilized with Cytofix/Cytoperm solution (BD) for 20 minutes at 4°C. Following one wash with Cytofix/Cytoperm wash buffer, cells were stained intracellularly with fluorochrome-conjugated antibodies specific for IFN-γ and TNF for 25 minutes at 4°C. Finally, cells were washed twice (once with Cytofix / Cytoperm wash buffer, once with FACS buffer) before resuspension in 200 μL of FACS buffer. All cells were run on a 5-laser Fortessa LSR II cytometer (BD) and analyzed in FlowJo (FlowJo LLC)

### In Vivo Cytotoxicity Assay

Adoptive transfer of fluorescently labeled, peptide-coated target splenocytes into LCMV-infected hosts for measurement of in vivo CD8+ T cell-mediated killing was completed as described previously (73). Briefly, spleens were collected from uninfected C57BL/6 mice and processed into a single-cell suspension as outlined above. Next, splenocytes were pulsed with HBSS or 1 µM of NP_396-404_ or GP_33-41_ peptide for 10 minutes at 37°C. Afterwards, the three populations of splenocytes were labeled with 2.5 µM, 1 µM or 0.4 µM of 5(6)-carboxyfluorescein diacetate succinimidyl ester (CFDA-SE; Invitrogen), which is metabolized within cells to carboxyfluorescein succinimidyl ester (CFSE), for 20 minutes at 37°C. Finally, labeled splenocytes were washed twice with Hank’s buffered saline solution and combined at an equal ratio. We then adoptively transferred ∼6×10^7^ splenocytes into each recipient mouse. After 4 hours, recipient mice were euthanized, and the survival of labeled populations was determined by flow cytometry. % lysis for each labeled population was calculated as follows: 100 – ([% LCMV target population in infected experimental/% unlabeled population in infected experimental) ÷ (% LCMV target population in naive control/% unlabeled population in naive control)] × 100).

### Complete blood count (CBC) analysis

Mice were anesthetized with continuous isoflurane and euthanized as described above. Complete blood counts were obtained with a Drew Scientific HemaVet 950 blood analyzer (Drew Scientific).

### Plasma ELISA analysis

Blood was collected as described previously. Following CBC analysis, blood was centrifuged to pellet RBCs. Plasma was aliquoted and frozen for further analysis. Plasma was thawed on ice at the time of sample preparation and diluted following manufacturer’s recommendations. ELISAs were performed as instructed by the manufacturing procedure (Insulin: Invitrogen; Cortisol: Arbor Assays; LDH: Sigma-Aldrich, PF4: Invitrogen). Plates were read by a BioTek Synergy H1 Plate reader (Agilent).

### Evans Blue analysis

On day 6 post-infection mice were intravenously injected with 150 µL/mouse of 2% Evans blue (Fisher Scientific, Waltham, WA). After two hours, mice were perfused with 10 mL PBS. Brains were weighed and incubated in 2 mL formamide at 55°C overnight. Tissues were vortexed briefly and centrifuged at 4,000 rpm for 10 minutes. 1 mL of supernatant was transferred to a plastic cuvette and fluorescence intensity was read at 620/680nm. An Evans blue standard curve was used to convert absorbance values to ng dye.

### Statistics

Statistical analyses were all performed using GraphPad Prism software. Each data set was first tested for normality following the Shapiro-Wilk test. If the data sets passed the normality test, a Student’s T-test or One-way ANOVA was performed depending on the number of data sets being analyzed. If the data was not found to be normal, then a Mann-Whittney or Kruskal-Wallis test was performed. For multiple comparisons, when statistical difference was determined, either a post-hoc Tukey or Dunn test was performed for normal and non-normal data, respectively.

### Study approval

All mouse protocols were compliant with the National Institutes of Health Guidelines for animal care and use. These studies were approved by Cincinnati Children’s Hospital Medical Center Institutional Animal Care and Use Committee and Institutional Biosafety Committee.

## Data availability

The authors declare that all supporting data and list of resources are available within the article. The data that support the findings of this study are available in the Supporting Data File.

## Acknowledgements

We thank Drs. Eric Mullins, Theodosia Kalfa, and Michael Jordan for their expertise and critical insights, and Ball Krishan Sharma, Laurel Romano, Benjamin Tourdot, and Jeremy Thompson for their technical assistance. Funding for this work was provided by NIH grants AR073228 (SNW), T32GM063483 (HAF), and 5T32AI118697 (RC). Support was also provided by the Cincinnati Children’s Research Foundation (SNW). The Flow Cytometry Core was supported by NIH grants AR070549 and DK078392.

## Author contributions

Conceptualization and design of study (RC, HAF, SNW, JSP), performance of experiments and data acquisition (RC, HAF, LR, AA, BG, CS), data analysis (RC, HAF, AA, DL, SNW, JSP), provision of reagents (JC, AR, BM), drafting of the manuscript (RC, HAF, SNW, JSP), and critical editing of the manuscript (RC, HAF, LR, AA, BG, CS, DL, JC, AR, BM, SNW, JSP). Authorship order among co-first authors was determined based on duration of involvement in this project.

## SUPPLEMENTAL MATERIAL

**Supplemental Figure 1.**
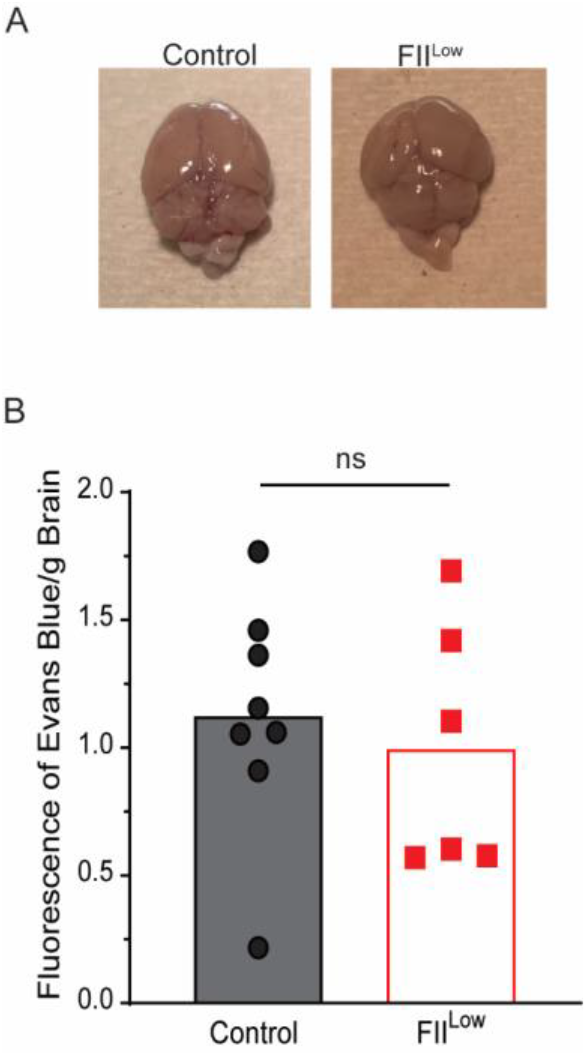
No evidence for vascular leakage in low prothrombin infected mice. C57BL/6 mice underwent pharmacologic depletion of prothrombin (FII^Low^, red squares) or control treatment (black circles) followed by intravenous infection with 2×10^6^ PFU of clone 13 LCMV (n=6-8/group). At day 6 of infection, Evan’s blue dye was administered intravenously two hours prior to euthanasia. Brains were harvested and analyzed (**A**) visually as well as (**B**) quantitatively. ns: p>0.05 by unpaired Student’s t-test.

**Supplemental Figure 2.**
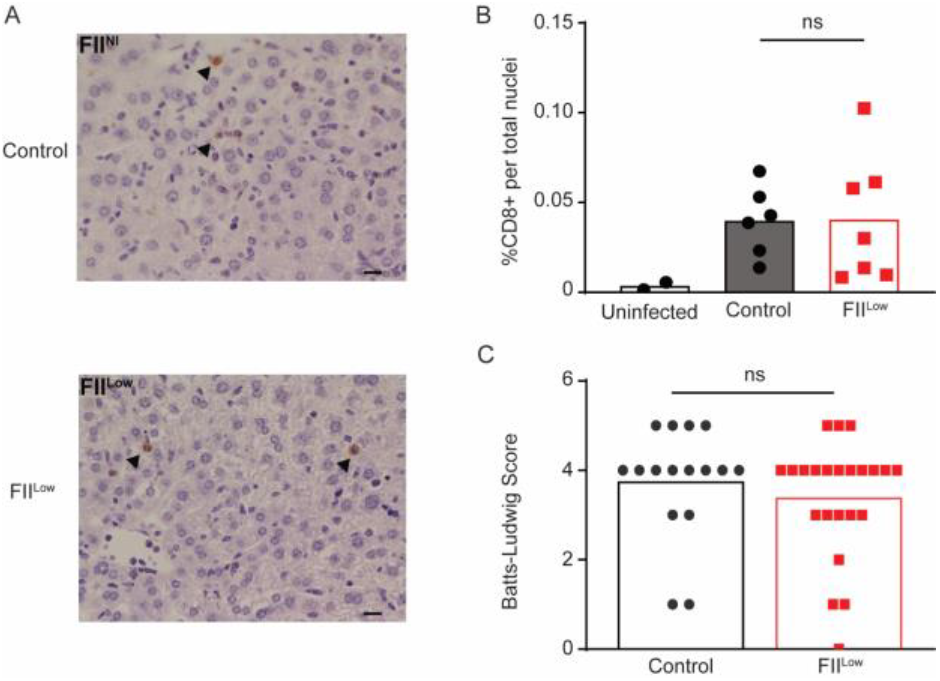
No enhanced liver infiltration of CD8 T cells in low prothrombin infected setting. C57BL/6 mice underwent pharmacologic depletion of prothrombin (FII^Low^, red squares) or control treatment (black circles) followed by intravenous infection with 2×10^6^ PFU of clone 13 LCMV (n=6-7/group, and 2 uninfected controls). At day 8 of infection, livers were harvested and stained for CD8α. (**A**) Representative images and (**B**) quantification are shown. (**C**) These livers sections and subjected to hematoxylin and eosin prior to blinded Batts-Ludwig scoring by a pathologist. Livers from uninfected mice (n=2) scored ≤1 in this assessment. ns: p>0.05 by unpaired Student’s t-test.

**Supplemental Figure 3.**
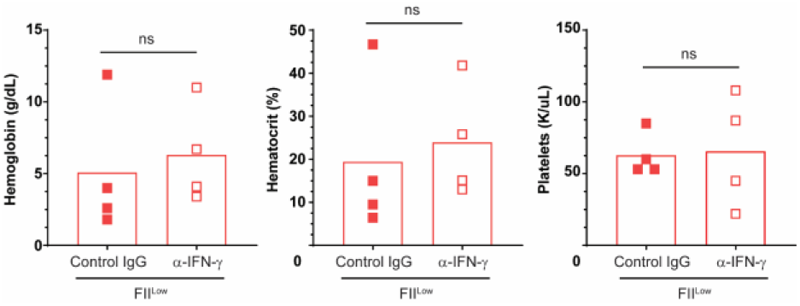
IFN-*γ* neutralization does not rescue anemia in infected low prothrombin mice. C57BL/6 mice underwent pharmacologic depletion of prothrombin (FII^Low^) followed by intravenous infection with 2×10^6^ PFU of clone 13 LCMV. On days 2 and 5 of infection, mice received retroorbital infections of 200 mg anti-mouse IFN-*γ* (α-IFN-*γ*, open red squares) or control IgG1 antibody (Control IgG, closed red squares, n=4/group). On day 8, blood was analyzed for hemoglobin levels, hematocrit, and platelet counts. ns refers to p>0.05 by unpaired Student’s t-test.

**Supplemental Figure 4.**
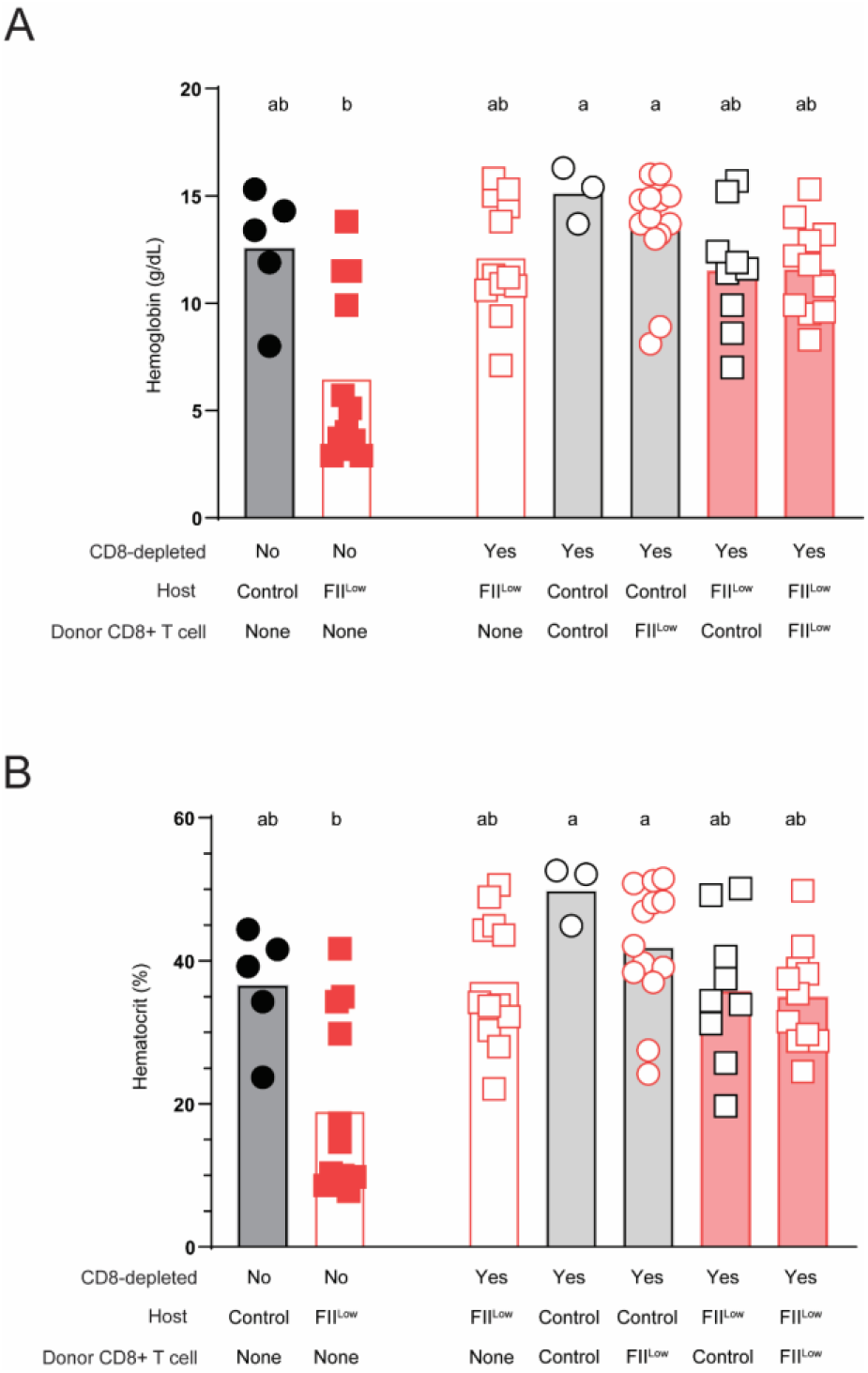
IFN-*γ* neutralization does not rescue anemia in infected low prothrombin mice. C57BL/6 mice underwent pharmacologic depletion of prothrombin (FII^Low^,) or control treatment. Some mice then received anti-CD8 antibodies (noted in top row as YES) to deplete CD8 T cells or isotype control antibody (Noted in top row as No) one day prior to intravenous infection with 2×10^6^ PFU of clone 13 LCMV. Donor mouse spleen CD8 T cells were isolated on day 6 and 100,000 of these cells (third denotes donor mouse source of CD8 T cells) were injected into groups of host mice (treatment listed on 2^nd^ row) on day 4 of infection. On day 8, recipient mouse blood was analyzed for (**A**) Hemoglobin and (**B**) hematocrit. Means followed by a common letter are not significantly different by ordinary one-way ANOVA, followed by Tukey’s multiple comparisons test, with single pooled variance.

